# Learning and attention increase visual response selectivity through distinct mechanisms

**DOI:** 10.1101/2021.01.31.429053

**Authors:** Jasper Poort, Katharina A. Wilmes, Antonin Blot, Angus Chadwick, Maneesh Sahani, Claudia Clopath, Thomas D. Mrsic-Flogel, Sonja B. Hofer, Adil G. Khan

**Author notes:** Correspondence to: JP or AGK.

## Abstract

Selectivity of cortical neurons for sensory stimuli can increase across days as animals learn their behavioral relevance, and across seconds when animals switch attention. While both phenomena are expressed in the same cortical circuit, it is unknown whether they rely on similar mechanisms. We imaged activity of the same neuronal populations in primary visual cortex as mice learned a visual discrimination task and subsequently performed an attention switching task. Selectivity changes due to learning and attention were uncorrelated in individual neurons. Selectivity increases after learning mainly arose from selective suppression of responses to one of the task relevant stimuli but from selective enhancement and suppression during attention. Learning and attention differentially affected interactions between excitatory and PV, SOM and VIP inhibitory cell classes. Circuit modelling revealed that cell class-specific top-down inputs best explained attentional modulation, while the reorganization of local functional connectivity accounted for learning related changes. Thus, distinct mechanisms underlie increased discriminability of relevant sensory stimuli across longer and shorter time scales.

## Introduction

Learning and attention both selectively enhance processing of behaviorally relevant stimuli (Gdalyahu et al., 2012; Goltstein et al., 2013; Li et al., 2008; McAdams and Maunsell, 1999; Ni et al., 2018; Reynolds and Chelazzi, 2004; Rutkowski and Weinberger, 2005; Schoups et al., 2001; Speed et al., 2020; Wiest et al., 2010; Yan et al., 2014; Yang and Maunsell, 2004). When animals learn what sensory features are task-relevant, or when they focus their attention on task-relevant features, early sensory cortical representations often undergo substantial changes. However, it is currently not known whether cortical changes during learning and attention rely on similar neural mechanisms.

The neural correlates of learning and attention share several characteristics. Visual learning results in increased stimulus selectivity through changes in stimulus-evoked neural firing rates (Gilbert and Li, 2012; Karmarkar and Dan, 2006; Li et al., 2008; Poort et al., 2015; Schoups et al., 2001; Yan et al., 2014; Yang and Maunsell, 2004), and is accompanied by changes in the interactions and correlations between neurons (Gu et al., 2011; Khan et al., 2018; Ni et al., 2018). Similarly, visual attention can also result in increased selectivity of attended stimuli, again through changes in stimulus-evoked firing rates (Reynolds and Chelazzi, 2004; Speed et al., 2020; Spitzer et al., 1988; Wimmer et al., 2015) and neuronal interactions (Cohen and Maunsell, 2009; Mitchell et al., 2009; Ni et al., 2018). Importantly, activity modulations during learning and attention are not uniformly distributed throughout the neural population but restricted to subsets of neurons (see for example (Chen et al., 2008; McAdams and Maunsell, 1999; Poort et al., 2015; Schoups et al., 2001; Yan et al., 2014)). Thus, both learning and attention lead to sharper and more distinct information being sent to downstream regions though subnetworks of learning- or attention-modulated cells.

Inhibition plays a crucial role in cortical plasticity (Froemke, 2015; van Versendaal and Levelt, 2016), and specific classes of inhibitory interneurons have been implicated in plasticity of cortical circuits during both learning and attention (Chen et al., 2015; Kato et al., 2015; Kuchibhotla et al., 2017; Makino and Komiyama, 2015; Sachidhanandam et al., 2016; Yazaki-Sugiyama et al., 2009). The activity of interneurons can change during both learning (Kato et al., 2015; Khan et al., 2018; Letzkus et al., 2011; Makino and Komiyama, 2015) and attention (Mitchell et al., 2007; Snyder et al., 2016; Speed et al., 2020), which can result in more stimulus-specific inhibition in the network.

Both learning and attention rely, to varying degrees, on the integration of top-down inputs with bottom-up signals. During attention, higher-order brain regions are thought to provide feedback signals to bias bottom-up information processing (Desimone and Duncan, 1995; Gilbert and Li, 2013), most prominently through direct feedback projections (Leinweber et al., 2017; Zhang et al., 2014) or through thalamic nuclei (Chalupa et al., 1976; Wimmer et al., 2015). These feedback projections can target excitatory or specific inhibitory interneurons (Leinweber et al., 2017; Zhang et al., 2014, 2016). In contrast, learning is thought to be primarily implemented by long-term plasticity of synapses, and reorganization of connectivity patterns (Froemke, 2015; Khan et al., 2018; Whitlock et al., 2006; Xiong et al., 2015), although top-down projections may also play a crucial role in guiding this process (Roelfsema and Holtmaat, 2018; Williams and Holtmaat, 2019).

Thus, both learning and attention modulate the firing properties of subsets of excitatory and inhibitory cortical neurons, leading to changes in firing rates and interactions between cells. It has therefore been suggested that learning and attention rely on similar neural mechanisms (Ni et al., 2018) or that attention-like processes may co-opt some of the underlying circuitry of learning (Kuchibhotla et al., 2017). However, this has never directly been tested, and it is not known if learning and attention engage the same neurons and circuits. A number of questions thus arise. First, within a population, is a common subset of neurons modulated by both learning and attention? Second, do learning-modulated and attention-modulated neurons undergo similar changes in their firing rates in order to increase stimulus selectivity? Third, do learning and attention result in similar changes in interactions between different excitatory and inhibitory cell classes?

To address these questions, we compared the changes in activity and interactions of the same population of neurons in V1 during learning and attention. We tracked the same identified pyramidal (PYR) neurons and parvalbumin (PV), somatostatin (SOM) and vasoactive intestinal peptide (VIP) positive interneurons as mice learnt to discriminate two visual stimuli and subsequently performed an attention switching task involving the same visual stimuli. We observed a similar profile of average changes in stimulus selectivity across the four cell classes during learning and attention. However, we discovered that these changes were largely uncorrelated at the single cell level, consistent with distinct mechanisms of selectivity changes during learning and attention. In support of this idea, we found that neural stimulus responses were dominated by selective suppression during learning, but displayed a combination of suppression and enhancement during attention. In addition, learning and attention differentially modulated interactions between excitatory and inhibitory cell classes. While learning-related changes were well captured by a model invoking changes in functional interaction strengths, attention-related changes were captured by a circuit model with top-down inputs targeted to PYR and SOM cells. These results reveal that more selective cortical representations for behaviorally relevant stimuli arise through distinct mechanisms over longer and shorter timescales.

## Results

### Increased response selectivity related to learning and attention switching

To understand how the same neural populations change their responses to visual stimuli with learning and attention, we trained mice to learn a go-no go visual discrimination task and subsequently trained them to perform an attention switching task involving the same pair of visual stimuli (Figure 1A,B). Head-fixed mice ran through a virtual approach corridor (Figure 1A) where the walls displayed a short stretch of circle patterns followed by grey walls for a random distance chosen from an exponential distribution (Figure 1C, top). Mice were then presented with one of two grating patterns, vertical or angled (40° relative to vertical), and were rewarded for licking a reward spout in response to the vertical grating. No punishment was given for licking the spout in response to angled gratings. All mice learned to discriminate the grating stimuli, reaching a threshold criterion of d′ > 2.0 (∼85% accuracy) within 7-9 days (Figure S1 example lick rasters from sessions pre- and post-learning. Figure 1D, average behavioral d-prime pre-learning -0.18 ± 0.56 s.d., post-learning 3.32 ± 0.82, sign test, P = 0.008, N = 8 mice).

**Figure 1.**
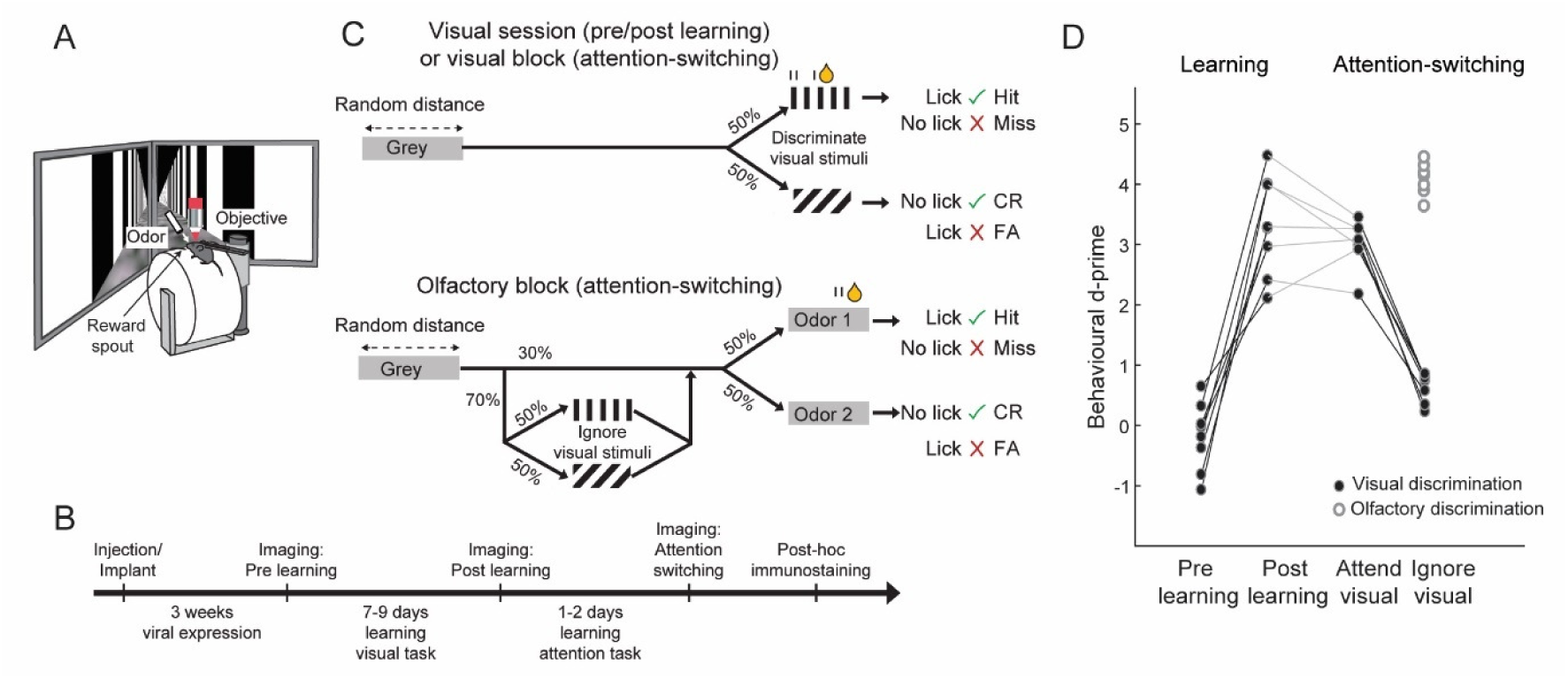
Visual discrimination learning and attention switching in mice. (A) Top, schematic showing virtual reality and imaging setup. (B) Experimental timeline. (C) Schematic of behavioral tasks. Top, visual discrimination: Mice were rewarded for licking the reward spout when vertical gratings were presented and not when angled gratings were presented. Olfactory discrimination: mice were rewarded for licking when odor 1 was presented and not when odor 2 or vertical or angled gratings were presented. (D) Behavioral discrimination performance (behavioral d’) across learning and during attention switching (N = 8 mice). Connected closed points indicate visual discrimination in individual mice. Open circles indicate olfactory discrimination.

We subsequently trained the mice to switch between blocks of the same visual discrimination task and an olfactory discrimination task, in which they learned to lick the reward spout to obtain a reward in response to one of two odors. During the olfactory discrimination blocks, the same grating stimuli used in the visual discrimination blocks were presented on 70% of trials but were irrelevant to the task (Figure 1C, bottom). Mice learnt this attention switching task in 1 to 2 days. Mice switched between the two blocks within the same session, successfully attending to and discriminating the grating stimuli in the visual block but ignoring the same grating stimuli while successfully discriminating odors during the olfactory blocks (Figure S1 example lick rasters from a session of attention switching behavior. Figure 1D, behavioral d-prime attend visual 3.02 ± 0.41 vs. ignore visual 0.63 ± 0.25, sign test P = 0.015, d-prime discriminating olfactory stimuli 4.10 ± 0.27).

### Selectivity changes at the population level are similar across learning and attention

We expressed the calcium indicator GCaMP6f in V1 using viral vectors and measured responses of L2/3 neurons using two-photon calcium imaging during the task. We re-identified the same neurons in co-registered, immunohistochemically stained brain sections from these animals and determined the identity of putative excitatory pyramidal (PYR) neurons and cells belonging to the three major classes of GABAergic inhibitory interneurons (Figure 2A). This approach allowed us to measure the simultaneous activity of PV, SOM and VIP positive interneurons along with the local excitatory neuron population (see Methods). We imaged the same 1249 PYR, 132 PV, 58 SOM and 175 VIP neurons before and after learning and a partially overlapping population of 5813 PYR, 477 PV, 245 SOM and 365 VIP neurons during the attention switching task (915, 105, 54 and 144 cells overlapping respectively, N = 8 mice).

**Figure 2.**
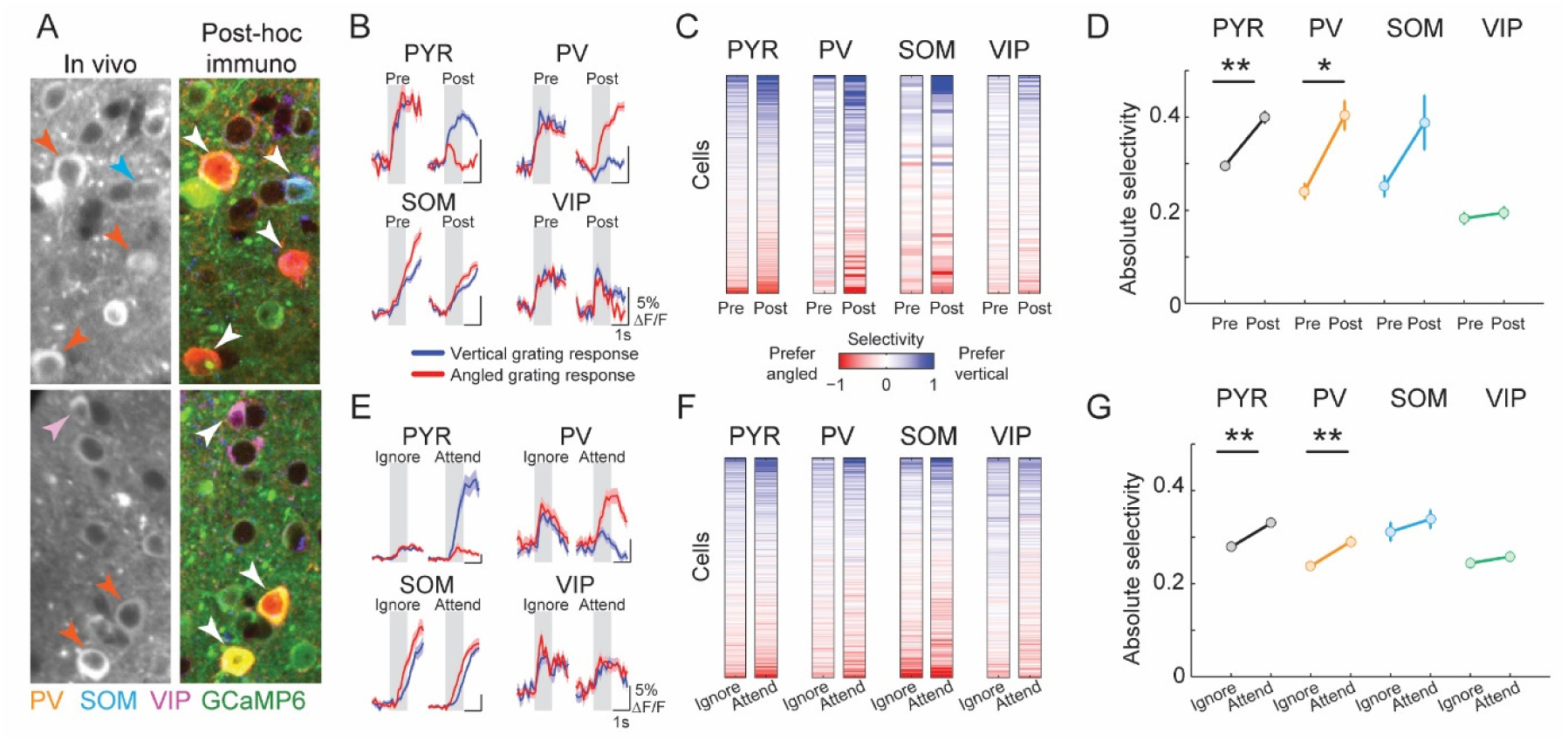
Similar changes in stimulus response selectivity across four cell classes during learning and attention switching. (A) Two example regions of in-vivo image planes with GCaMP6f-expressing neurons and the same regions after post hoc immunostaining for PV, SOM and VIP (orange, blue and magenta, respectively) following image registration. Identified interneurons are indicated by arrowheads. (B) Example cells from the 4 cell classes, average responses to vertical (blue line) and angled (red line) grating stimuli before (pre) and after (post) learning. Shaded area represents SEM. Gray shading indicates 0-1s window from stimulus onset used to calculate stimulus selectivity. (C) Stimulus selectivity of the same cells (rows) before and after learning (columns). Cells were ordered by their mean pre- and post-learning selectivity. Numbers of cells recorded both pre- and post-learning: 1,249 PYR, 132 PV, 58 SOM and 175 VIP cells, here and in D. (D) Average absolute selectivity of the 4 cell classes before and after learning. Error bars represent SEM. Sign test, **P < 0.001; *P < 0.05. Selectivity distribution in Figure S4A. (E-G), Same as B-D for attention switching task. Numbers of cells recorded: 5813 PYR, 477 PV, 245 SOM and 365 VIP cells.

Neurons from each cell class showed varying degrees of responsiveness to the visual grating stimuli (Figure S2A,B). During learning, we observed changes in visual grating responses in subsets of neurons from all cell classes (Figure 2B, Figure S2A,B). This led to changes in stimulus selectivity (difference in the mean responses to the two grating stimuli normalized by response variability, see Methods) in individual cells to varying degrees (Figure 2C). On average, PYR and PV cells significantly increased their stimulus selectivity during learning, as reported previously (Khan et al., 2018; Poort et al., 2015) (Figure 2D; PYR, average absolute selectivity pre-learning, 0.30 ± 0.31 (mean ± s.d.), post-learning 0.40 ± 0.44, sign test, P < 10^−9^, N = 1249, PV, pre-learning, 0.24 ± 0.19, post-learning 0.40 ± 0.36, P = 0.002, N = 132). In contrast, the average selectivity of SOM and VIP interneurons did not change significantly (SOM, pre-learning 0.25 ± 0.17, post-learning 0.39 ± 0.45, P = 0.51, N = 58, VIP, pre-learning 0.18 ± 0.16, post-learning 0.20 ± 0.17, P = 0.45, N = 175).

We found a similar profile of selectivity changes across cell classes between the ‘ignore’ and ‘attend’ conditions of the attention switching task. Specifically, visual stimulus selectivity increased on average in PYR and PV cells but not in SOM and VIP cells when mice switched from ignoring to attending the same visual grating stimuli (Figure 2E-G; PYR, ignore 0.28 ± 0.28, attend 0.33 ± 0.32, P < 10^−10^, N = 5813, PV, ignore 0.24 ± 0.18, attend 0.29 ± 0.25, P = 0.0007, N = 477, SOM, ignore 0.31 ± 0.31, attend 0.34 ± 0.31, P = 0.25, N = 245, VIP, ignore 0.24 ± 0.19, attend 0.26 ± 0.19, P = 0.60, N = 365). Changes in running and licking could not account for the increased selectivity of responses during learning or attention (Figure S3A,B). Thus, learning and attention both led to similar changes in stimulus selectivity of V1 neurons on average, across excitatory and multiple inhibitory cell classes.

### Selectivity changes at single cell level are uncorrelated

The similar profile of changes in average selectivity during learning and attention switching suggested that the neural basis of these two changes may be overlapping. Indeed, both learning and attention serve a similar purpose: to enhance an animal’s ability to detect and respond to relevant stimuli, and prior work has suggested that the two may be implemented by common neural mechanisms (Ni et al., 2018). We therefore asked whether the increase in selectivity during learning and attention was related at the single neuron level.

Across the population of PYR neurons, we found that there was no significant correlation between the learning related and attention related changes in stimulus selectivity (Figure 3A, R = 0.01, P = 0.67, see also Figure S2C). This indicated that a cell’s change in stimulus selectivity during learning had no bearing on its change during attention. This absence of correlation was not due to extensive changes in the original visual response selectivity of these cells from the post-learning session to the attention switching session – there was a strong correlation between the post-learning selectivity and the selectivity during the attend condition of the attention switching task (Figure 3B, R = 0.61, P < 10^−99^).

**Figure 3.**
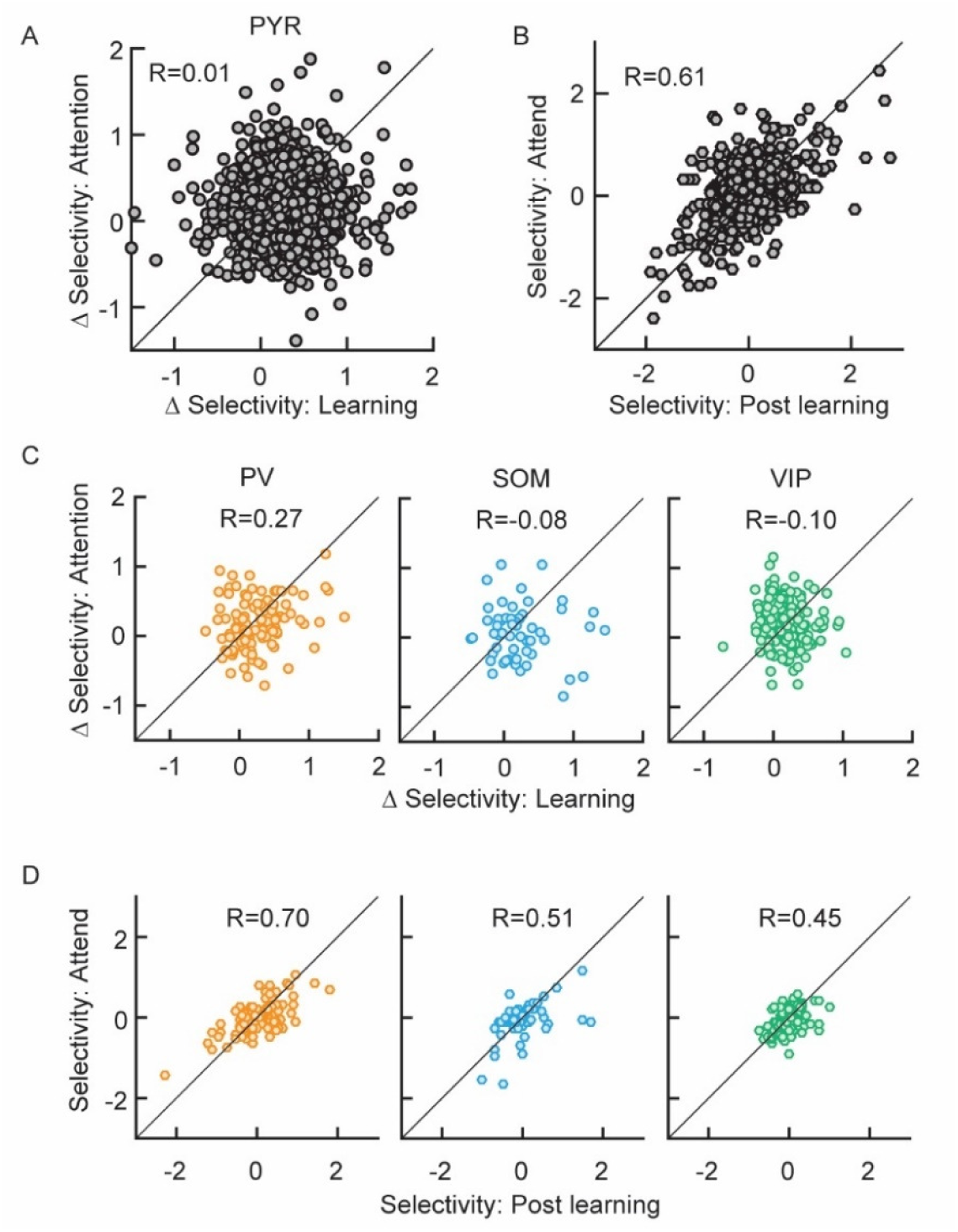
Changes in stimulus selectivity during learning and attention are uncorrelated. A) Relationship between ΔSelectivity with learning (positive values indicate increased selectivity after learning) and ΔSelectivity with attention (positive values indicate increased selectivity with attention) for PYR cells (N = 915 cells). B) Relationship between post-learning selectivity and selectivity in the attend condition for PYR cells. C, D) Same as A and B for the three interneuron classes (N = 105 PV, 54 SOM and 144 VIP cells).

We observed a moderate but significant correlation between the learning-related and attention-related changes in stimulus selectivity in PV interneurons, but not SOM or VIP interneurons (Figure 3C, PV, R = 0.27, P = 0.01, SOM, R = 0.08, P = 0.57, VIP, R = 0.10, P = 0.25), raising

the possibility that subsets of PV cells may be preferentially engaged in both learning and attention. All interneuron cell classes displayed strong correlations between the post-learning selectivity and the selectivity during the attend condition (Figure 3D, PV, R = 0.70, P < 10^−16^, SOM, R = 0.51, P < 10^−4^, VIP, R = 0.45, P < 10^−8^), again ruling out extensive changes in the stimulus tuning of cells between the post-learning and attention switching sessions.

Thus, while increases in neural selectivity due to learning and attention were similar across excitatory and multiple inhibitory interneuron classes on average, they were largely uncorrelated at the single cell level. The lack of correlation between selectivity modulations during learning and attention suggested that these two processes may be driven by distinct neural mechanisms.

### Mechanisms of selectivity change

Neurons can increase their stimulus selectivity by selective suppression of responses to non-preferred stimuli (Lee et al., 2012), selective increase in responses to preferred stimuli (McAdams and Maunsell, 1999) or a combination of the two. We tested for the relative prevalence of these changes in the population of PYR cells during learning and attention.

First, we studied changes in stimulus-evoked firing rates in all recorded PYR cells, regardless of their stimulus selectivity. We subtracted the pre-learning from the post-learning stimulus response profile of each cell for a given stimulus, to obtain the difference-PSTH. During learning, the difference-PSTHs of the PYR population were dominated by cells with negative deflections from baseline, i.e. cells which decreased their stimulus response amplitude to the same stimulus during learning (Figure 4A, left). This was true for both rewarded and non-rewarded stimuli (Figure S5A, left). Interestingly, the difference-PSTH during attention switching (attend minus ignore condition), revealed that changes with attention were more uniformly distributed across increases and decreases in response amplitude (Figure 4A, right). This was again true for both rewarded and non-rewarded stimuli (Figure S5A, right, difference-PSTH averaged 0-1s significantly different between learning and attention, P < 10^−28^, sign test, Figure S5D). Thus, learning, unlike attention, was dominated by a suppression of responses.

**Figure 4.**
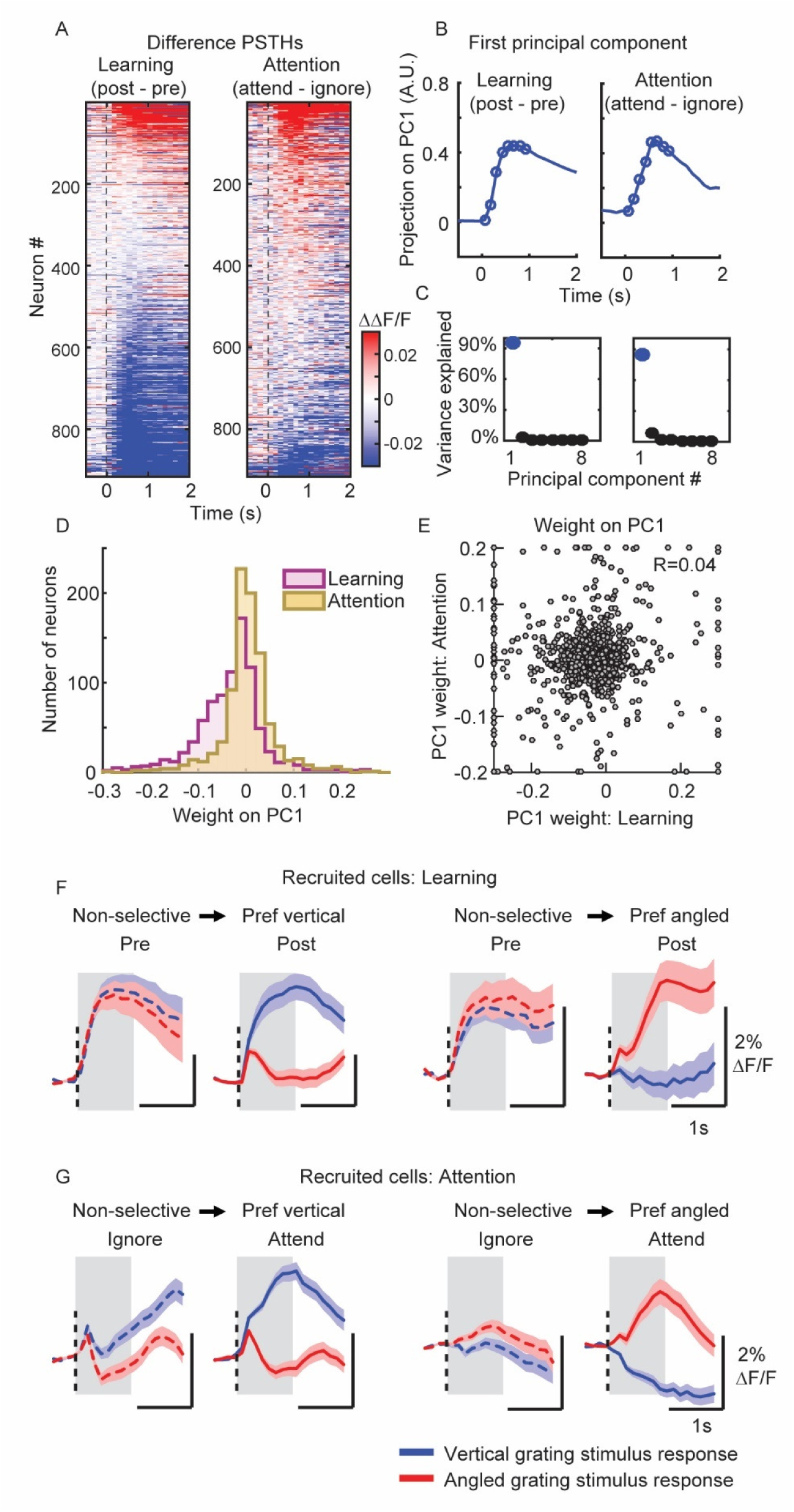
Increased stimulus selectivity through selective response suppression during learning but enhancement and suppression during attention. A) Difference in calcium responses to the rewarded vertical grating stimulus, post minus pre learning (left) or attend minus ignore conditions (right) for all recorded PYR cells (Difference-PSTHs). Responses are baseline corrected (subtraction of baseline ΔF/F –0.5 to 0 s before stimulus onset) and aligned to grating onset (dashed line). Cells are sorted by their average amplitude 0–1 s from stimulus onset. N = 915 matched cells, in A to E, N = 8 mice. B) First principal component (PC) of the difference-PSTHs from the learning (left) and attention data (right). Circles indicate the time points (0-1s) used to determine the PCs. C) Percentage of variance explained by each PC during learning (left) and attention (right). D) Distribution of weights from each cell onto the first PC during learning and attention. E) Relationship between the weights of cells on the first PC during learning and attention. Values greater than the axis limits are pegged to the maximum displayed value. F) Average PSTHs of all recruited cells, i.e. cells which changed from non-selective to selective stimulus responses during learning, N = 243 and 216 cells recruited with preference for vertical stimulus or angled stimulus respectively. G) Average PSTHs of all recruited cells during attention, N = 672 and 676 cells recruited with preference for vertical stimulus or angled stimulus respectively. Shaded area represents SEM. Gray shading indicates 0-1s window from stimulus onset used for analysis.

Learning and attention might lead to complex temporal changes in firing rate profiles, not captured in the above analysis. We therefore performed principal component analysis (PCA) to identify the components which captured the majority of variance in the shapes of all difference-PSTHs. Interestingly, for both learning and attention, we found that a single component accounted for more than 80% of the variance across all cells, and this component was highly similar for both learning and attention (Figure 4B, C). However, the distributions of weights projected onto this PC during learning and attention were substantially different, with a predominance of negative weights during learning (Figure 4D, P < 10^−38^, sign test). Thus, while we did not find a difference in the temporal profile of firing rate changes, we confirmed the robust presence of stimulus response suppression during learning, but not during attention.

At the single cell level, we found that the scores on the first PCA components were uncorrelated (Figure 4E, R = 0.04, P = 0.24, see Figure S5E for a similar effect with average calcium responses), suggesting independent firing rate modulation of individual cells by learning and attention.

We next asked what changes in firing rates underlie the increased stimulus selectivity in the population. We restricted this analysis to recruited cells, that is, cells which changed from non-selective to significantly selective during learning or attention. The average PSTHs of these cells showed markedly distinct features. During learning, recruited cells showed preferential suppression of responses to one of the two stimuli (Figure 4F). In contrast, with attention, cells became selective through a combination of enhancement and suppression of responses to the two stimuli (Figure 4G). (Percent changes in stimulus response amplitude to vertical and angled stimuli: Figure 4F left, -8%, -81%, Figure 4F right -89%, -27%. Figure 4G left, 72%, 10% (not significant), Figure 4G right -92%, 56%. Changes calculated as the percentage of the maximum in each category, all responses averaged 0-1s, all P values < 10^−4^ except where stated).

Thus, learning was associated with suppression of evoked responses, particularly of the non-preferred stimulus, while attention was mainly associated with increased responses of the preferred stimulus.

### Changes in interactions between excitatory and inhibitory cell classes

Changes in cortical processing are accompanied by a reconfiguration of network dynamics and interactions. We previously demonstrated that interactions between PV cells and surrounding PYR cells are reorganized during learning (Khan et al., 2018). Specifically, we measured the correlation between PV cell selectivity and the selectivity of the PYR cell population within 100 μm of each PV cell. The slope and correlation coefficient of this relationship significantly decreased during learning (Figure 5A top, pre learning, slope = 0.21, confidence intervals (CI) 0.14 to 0.29, R = 0.48, post learning, slope = 0.05, CI 0.00 to 0.10, R = 0.19, reduction in slope bootstrap test P < 10^−4^), suggesting that during learning, PV cell activity became less dependent on the average stimulus preference of surrounding PYR cells. However, when we performed the same analysis comparing ignore and attend conditions, we found no difference in the correlation coefficient or slope of this relationship (Figure 5A bottom, ignore, slope = 0.05, CI 0.03 to 0.07, R = 0.23, attend, slope = 0.03, CI 0.01 to 0.05, R = 0.15, reduction in slope bootstrap test P = 0.06). Indeed, the relationship appeared similar to that observed at the end of learning. This was despite the fact that PV cells displayed a comparable degree of selectivity increase with attention as with learning.

**Figure 5.**
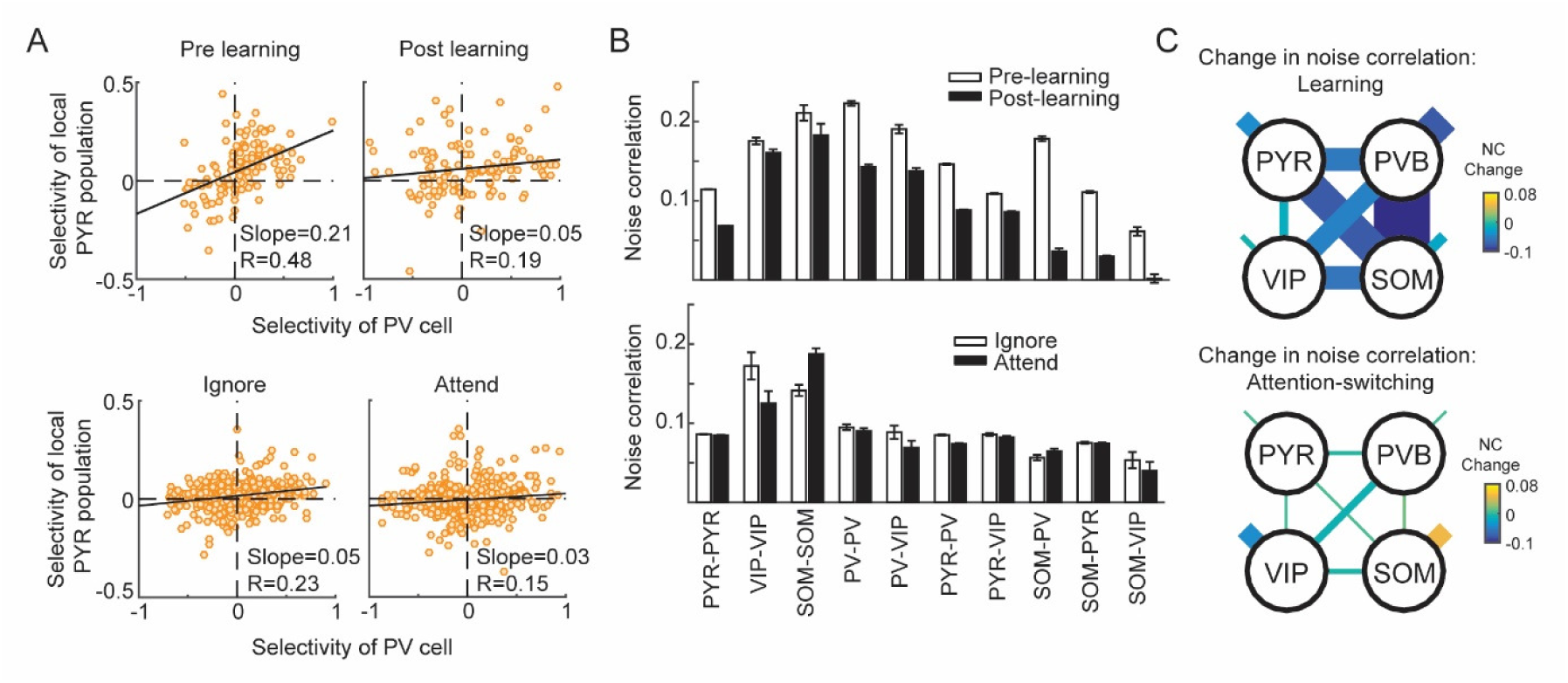
Distinct changes in interactions between excitatory and inhibitory cells during learning and attention. A) Top, relationship between the selectivity of individual PV cells and the mean selectivity of the local PYR population within 100 μm of each PV cell, before (pre) and after learning (post). Bottom, same comparison for the ignore and attend conditions of the attention switching task. B) Average noise correlations between cell pairs belonging to the same or different cell classes, before and after learning (top) or in the ignore and attend conditions (bottom). Only cells with significant responses to the grating stimuli were included. The number of cell pairs in each cell class combination was as follows: pre-, post-learning, PYR–PYR 74,581, 64,921; VIP–VIP 1166, 907; SOM–SOM 215, 99; PV– PV 1,731, 1,369; PV–VIP 790, 718; PV–PYR 17,792, 15,283; PYR–VIP 14,681, 12,009; SOM–PV 1,250, 690; SOM–PYR 7,112, 4,952; SOM–VIP 455, 377. Ignore/attend conditions, PYR–PYR 61,175; VIP–VIP 58; SOM–SOM 381; PV–PV 777; PV–VIP 129; PV–PYR 11,312; PYR–VIP 3024; SOM–PV 814; SOM–PYR 6,626; SOM–VIP 136. Error bars represent SEM. Full data distribution can be seen in Figure S4B. C) Changes in noise correlations (shown in B) due to learning (top) or attention (bottom) as indicated by line thickness and color code. Shorter line segments indicate change in noise correlations between cells of the same type.

To further explore the network signatures of changes during learning and attention, we computed noise correlations during the grating stimulus period between pairs of neurons within and across cell classes, before and after learning and during attend and ignore conditions. Since noise correlations are a measure of the stimulus-independent trial-to-trial co-variability of neural responses, they provide an estimate of mutual connectivity and shared inputs. As reported earlier, we found that during learning, SOM cells become de-correlated from pyramidal, PV and VIP neurons, with the largest changes between cell classes (sign test, all reductions in noise correlation were significant, with the exception of SOM–SOM cell pairs, P=0.8, see also (Khan et al., 2018)). Specifically, we observed a large reduction in noise correlation between SOM-PV, SOM-PYR and SOM-VIP cell pairs during learning (Figure 5B,C, top, vertical grating stimulus. Full distributions in Figure S4B).

In contrast, during attention switching, we found that the largest absolute changes in noise correlation were within cell classes, namely between SOM-SOM and VIP-VIP cell pairs (Figure 5B,C bottom). SOM-SOM cell pairs displayed an increase in noise correlation (sign test, P < 10^−9^) whereas VIP-VIP pairs displayed decreased noise correlation (P = 0.02). In addition, PYR-PYR, PYR-PV, PYR-SOM and PV-PV cell pairs also showed a significant reduction in noise correlation, although the absolute change was smaller (all Ps < 0.03). Changes in running speed or licking could not account for the observed changes in noise correlations (Figure S3C,D).

Thus, learning and attention are associated with different patterns of changes in noise correlations between excitatory and multiple inhibitory cell classes, consistent with the idea that distinct mechanisms underlie these processes.

### Modelling response changes during learning and attention

What changes in network properties underlie the observed changes during learning and attention? We recently developed a multivariate autoregressive (MVAR) linear dynamical system model to predict the activity of single cells based on interaction weights with their local neighbors. Analysis of the MVAR model fit to the neural responses during learning revealed that increased response selectivity after learning was associated with the reorganization of interaction weights between cells (Figure S6A-C see also (Khan et al., 2018)). We tested if similar changes in functional connectivity can account for the changes in stimulus responses observed with attention. We compared a model that allowed interaction weights to change across the attend and ignore conditions against a simpler model that used the same weights across both conditions. We found that the fit quality of the MVAR model, quantified by the cross-validated R^2^, was actually lower for the model allowing weights to change across the attend and ignore conditions, demonstrating that changing interaction weights during attention conferred no advantage to the model (Figure S6B). Even when weights were allowed to change in the MVAR model, we found stable PYR-PV interaction weights, in contrast to the changes in weights observed during learning (Figure S6C). Together with the absence of reorganization of PYR-PV interactions during attention (Figure 5A, bottom), these results suggest that functional connectivity is relatively stable during attention, but changes during learning, possibly through long-term synaptic plasticity mechanisms.

Since the data-driven MVAR model analysis indicated that the selectivity changes were not predicted by changes in local functional interactions, we developed a detailed theoretical model of the local circuit enabling us to evaluate what type of external inputs could explain the attentional modulation of the local circuit. In this model, we represented each of the four cell types (PYR, PV, SOM, VIP) by their population activity, corresponding to the average response across all cells with a given stimulus preference in the population. Population activity was determined by baseline activity, feedforward stimulus-related input, top-down attentional modulatory input, and connection weights with other cell populations (see Methods). The four neural populations were connected using experimentally derived connectivity values, similar to (Kuchibhotla et al., 2017) (Figure 6A). The model’s population responses resembled the average population stimulus responses of all four cell classes (Figure 6B) (Khan et al., 2018).

**Figure 6.**
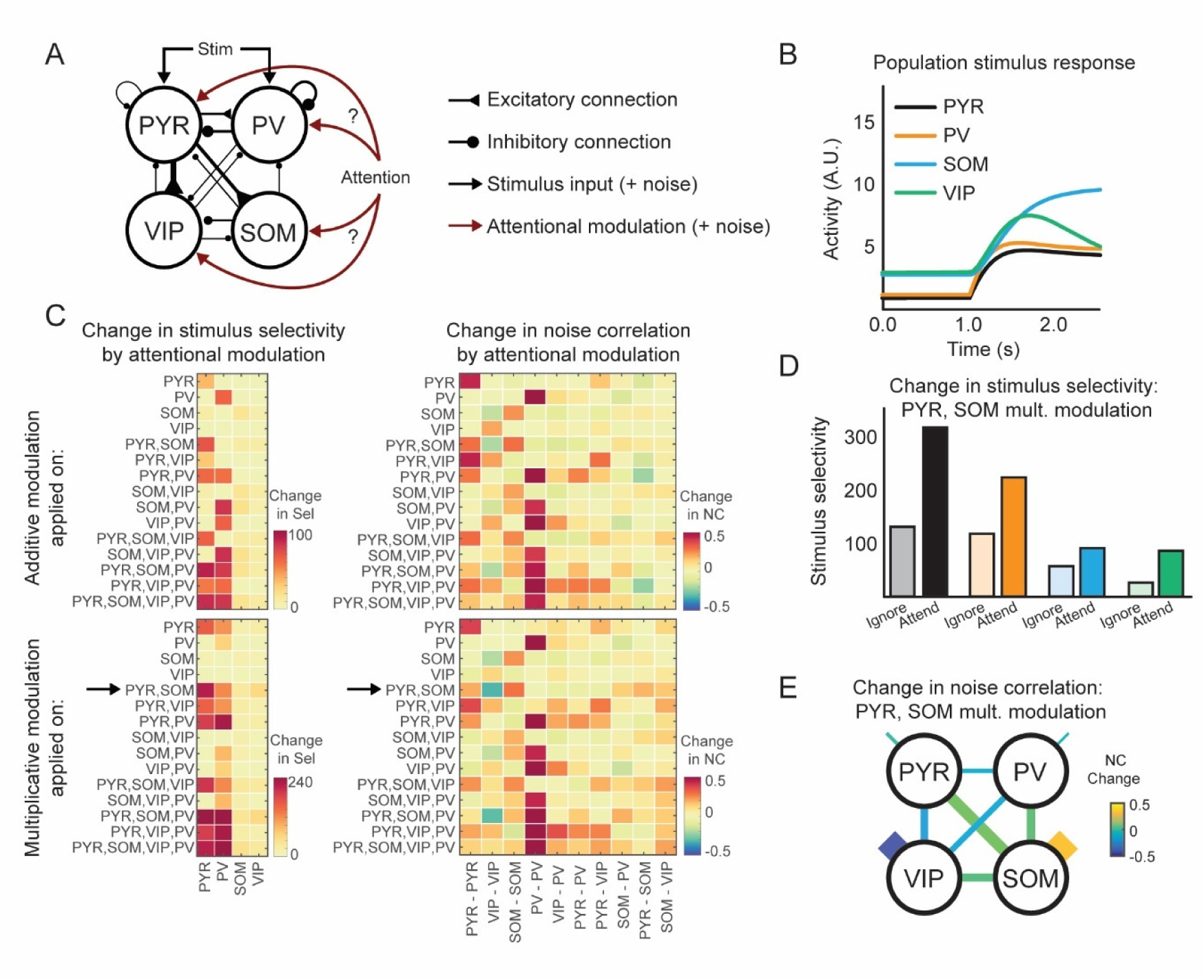
A circuit model can distinguish between different patterns of top-down attentional modulation. (A) The model architecture, indicating connectivity between different cell classes and possible sources of shared external fluctuations. (B) Simulated responses of the four cell types to the preferred stimulus. (C) Changes in stimulus selectivity and noise correlations (NC) obtained from models with attentional modulation applied to different combinations of cell populations. Both additive and multiplicative modulations were tested. Arrow indicates the condition which best replicated the experimental changes in selectivity and noise correlation. (D) Absolute selectivity of different cell classes without (Ignore) and with (Attend) attentional modulation provided to PYR and SOM populations, with PYR receiving 0.7 times the modulation of SOM (see Figure S6D,E). (E) Changes in noise correlations (NC change) with attentional modulation as in (D) between and within the four cell classes, as indicated by line thickness and color code.

In the model, each population received fluctuations from cell-intrinsic sources (e.g. due to ion channel noise) and shared external sources (stimulus and top-down modulatory inputs, Figure 6A). The simulated noise correlations thus reflected both connectivity and fluctuations in the stimulus and modulatory inputs. Since functional connectivity weights between cell classes were stable across attend and ignore conditions, we modelled the changes in noise correlations during attention switching as arising from changes in the shared external fluctuations.

It is unclear whether attention has a multiplicative effect (Goris et al., 2014; Reynolds and Heeger, 2009) or an additive effect (Buracas and Boynton, 2007; Thiele et al., 2009). We therefore considered two different types of models with an additive or multiplicative effect of attentional modulation. We systematically simulated all conditions in which attentional modulation targeted different cell classes and combinations of cell classes. We then evaluated the stimulus selectivity changes and noise correlation changes induced by attentional modulation (Figure 6C). We looked for conditions which replicated our experimental findings, including (a) attention increased only PYR and PV stimulus selectivity (Figure 2G) and (b) attention mainly increased SOM-SOM and decreased VIP-VIP noise correlations (Figure 5C, bottom). Of all conditions, only one matched both these experimental findings, where PYR and SOM cells received multiplicative attentional modulation (Figure 6C, arrows).

The model so far assumed equal influence of attentional modulation onto all cells. We next varied the relative strengths of modulation received by PYR and SOM cells to test whether the match to experimental findings could be improved. Specifically, the current model produced an increase in noise correlations between PYR-PYR, PYR-SOM, SOM-PV and SOM-VIP cells, which was not observed experimentally. A model in which the attentional modulation of PYR was 0.7 times the modulation of SOM improved the match to the data (Figure S6D). This model replicated the increase in PYR and PV stimulus selectivity (Figure 6D) as well as the changes in SOM-SOM and VIP-VIP noise correlations, with only minor changes in noise correlations between other cell types (Figure 6E). Thus, a model in which PYR and SOM populations received different degrees of multiplicative attentional modulation best accounted for the changes in selectivity and noise correlations observed in the data (Figure S6E).

## Discussion

We show that improvements in sensory coding arising from learning or attention rely on distinct mechanisms, based on three lines of evidence. First, at the single-cell level, the effects of learning and attention are uncorrelated. Second, distinct firing rate changes underlie the increases in selectivity during learning and attention. Third, learning and attention are associated with different changes in functional interactions between cell classes. Our computational models suggest that learning relies on reorganization of interactions in the local circuit, whereas attention relies on multiplicative top-down signals that target specific cell-classes.

### Subpopulations of excitatory neurons modulated by learning and attention

Learning and attention are closely linked: attended objects are preferentially learnt, and learning can bias the allocation of attention (Gilbert et al., 2000; Vartak et al., 2017). Although we show that learning and attention both lead to a similar increase in stimulus selectivity on average in PYR cells, these increases are not driven by the same subset of neurons. Importantly, this does not mean that cells are either modulated by learning or attention. Instead, learning and attention each modulate the same neurons to varying degrees, and a neuron’s degree of modulation during learning is uncorrelated with its degree of modulation by attention.

The basis of neural susceptibility to either learning- or attention-related modulations is poorly understood. For example, it may be related to intrinsic excitability (Brebner et al., 2020), expression of immediate-early genes (e.g. CREB (Han et al., 2007) or Arc (Gouty-Colomer et al., 2016), see also (Holtmaat and Caroni, 2016)), and pre- or post-synaptic expression of neuromodulator receptors (Disney et al., 2007; Herrero et al., 2008), or connectivity with distal and top-down inputs (Iacaruso et al., 2017; Marques et al., 2018). Our results impose an important restriction: these molecular or circuit mechanisms must be independent or exert a minimal influence on each other, since the effects of learning and attention on individual pyramidal cells are uncorrelated.

We found a small but significant correlation in the learning- and attention-related selectivity changes in PV interneurons. Given the lack of correlation for PYR cells, and the fact that this effect is difficult to account for by a model in which PV cells inherit their stimulus response properties from neighboring PYR cells (Kerlin et al., 2010; Khan et al., 2018; Scholl et al., 2015) this effect requires further investigation.

### Suppression and enhancement of stimulus responses

We find that learning and attention lead to distinct patterns of suppression and enhancement of firing rates. Learning was dominated by selective suppression of responses to the non-preferred stimulus, perhaps because it is metabolically more efficient for implementing long-term selectivity changes (Howarth et al., 2012). Previous studies of associative conditioning have described both suppression and enhancement of responses in sensory cortex (Gdalyahu et al., 2012; Goltstein et al., 2013; Makino and Komiyama, 2015). By longitudinally tracking the same neurons, we find that learning is largely accompanied by sparsification of cortical responses. Attention, in contrast, largely led to selectivity changes through selective enhancement of responses. This is consistent with a large body of work showing that enhancement of attended responses is a common form of attentional modulation (McAdams and Maunsell, 1999; Speed et al., 2020; Spitzer et al., 1988; Wilson et al., 2019). Here, by studying the same neural population across both learning and attention, we demonstrate that V1 neurons are remarkably versatile, capable of displaying either selective enhancement or selective suppression of stimulus responses according to the current behavioural demand.

### Changes in interactions

Imaging the activity of multiple cell classes simultaneously allowed us to investigate both interactions within and between excitatory and inhibitory cell classes. We found changes in interactions at two levels.

First, we observed a reorganization of interaction weights between PYR and PV cells during learning, possibly through long-term synaptic plasticity, which was captured quantitatively by a linear dynamical systems model. In contrast, attention did not lead to a similar change in interaction weights, suggesting that the short timescale of attention does not permit large-scale reorganization of connectivity patterns.

Second, we found changes in noise correlations between pairs of the same or different cell classes. Changes in noise correlations have been implicated in improved behavioral abilities during learning and attention (Jeanne et al., 2013; Ni et al., 2018). We found that noise correlation changes were dramatically different across learning and attention. Learning was marked by reductions in inter-cell class correlations. Specifically, SOM cells became decorrelated from the rest of the network. This transition potentially facilitates plasticity in the network, by reducing the amount of dendritic inhibition from SOM cells that coincides with visual responses in excitatory cells (Khan et al., 2018). In contrast, attention changed correlations of SOM-SOM and VIP-VIP cell pairs, leaving inter cell-class correlations relatively unchanged. Our model demonstrates that these changes can be explained by top-down input in the absence of local connectivity changes. Importantly, this relies on specific connectivity motifs across cell classes (Fino and Yuste, 2011; Hofer et al., 2011; Jiang et al., 2015; Pfeffer et al., 2013).

To account for the increased stimulus selectivity and noise correlation changes, we tested a variety of circuit architectures (Prinz et al., 2004). Top-down attentional modulation signals can be multiplicative (Goris et al., 2014; Reynolds and Heeger, 2009) or additive (Buracas and Boynton, 2007; Thiele et al., 2009), and they can target specific cell classes (Leinweber et al., 2017; Zhang et al., 2014, 2016). Here, the experimental results limited possible model architectures to a single one, with multiplicative top-down modulation targeting SOM and PYR cells. Top-down projections with specific targeting have been proposed to be central to the gating of plasticity, allowing attention to guide learning (Roelfsema and Holtmaat, 2018). These specific predictions of targeted top-down projections provide a basis for future experimental work.

In summary, learning and attention lead to similar increases in neural response selectivity, but the effects are driven by different subsets of cells. Cells undergo distinct patterns of activity changes to achieve increased neural response selectivity during learning and attention. These results highlight the remarkable versatility by which a cortical circuit implements computations across short and long time scales.

## Acknowledgements

We thank the GENIE Program and Janelia Research Campus of the Howard Hughes Medical Institute for making GCaMP6 material available. This work was supported by the European Research Council (SBH, HigherVision 337797; TDM-F, NeuroV1sion 616509), the SNSF (SBH, 31003A_169525, AGK, PZ00P3_168046), EMBO (AB, ALTF 74-2014), the Wellcome Trust (AGK 206222/Z/17/Z, JP 211258/Z/18/Z, CC 200790/Z/16/Z, TDM-F & SBH 090843/F/09/Z), the BBSRC (CC, BB/N013956/1, BB/N019008/1), the EPSRC (CC, EP/R035806/1), the Simons Foundation (CC 564408, MS, SCGB 323228, 543039), the Gatsby Charitable Foundation (MS, TDM-F, SBH GAT3361), the DFG (KAW 398005926) and Biozentrum core funds (University of Basel).

## Author contributions

JP, TDM-F, SBH and AGK designed the experiments. JP and AGK performed the experiments and analyzed the data. KW developed and analyzed the circuit model with supervision from CC. AC developed and analyzed the MVAR model with supervision from MS. AB performed the immunostaining and contributed to the post hoc cell matching procedure. All authors discussed the data. JP and AGK wrote the paper, with inputs from all authors.

## Lead Contact

Further information and requests for resources and reagents should be directed to and will be fulfilled by the lead contacts and corresponding authors Jasper Poort and Adil Khan.

## Materials Availability

This study did not generate new unique reagents

## Data and code availability

The data and code that support the findings of this study are available from the corresponding authors upon request.

## Methods

Experimental procedures for the behavioral task, surgery, two-photon calcium imaging, post-hoc immunostaining and image registration have been described in detail in previous studies (Khan et al., 2018; Poort et al., 2015).

### Animals and two-photon calcium imaging

All experimental procedures were carried out in accordance with institutional animal welfare guidelines and licensed by the UK Home Office and the Swiss cantonal veterinary office. Mice were C57Bl/6 wild type mice (3 males, 1 female, Janvier Labs), crosses between Rosa-CAG-LSL-tdTomato (JAX: 007914) and PV-Cre (JAX: 008069) (2 males), and crosses between Rosa-CAG-LSL-tdTomato and VIP-Cre (JAX: 010908) (1 male, 1 female) all obtained from Jackson Laboratory. Data from these mice were used in a prior study (Khan et al., 2018).

Mice aged P48-P58 were implanted with a chronic imaging window following viral injections of AAV2.1-syn-GCaMP6f-WPRE (Chen et al., 2013). Multi-plane two-photon imaging began approximately three weeks after surgery, during which 4 planes were imaged with 20 µm spacing at an imaging rate of 8 Hz for each imaging plane. Eight mice were imaged both pre-learning (either first or second day of training) and post-learning (either day 7, 8 or 9 of training), and during an attention switching task (1 session each, after 1 to 2 days of learning the attention switching task). Before each imaging session the same site was found by matching anatomical landmarks.

### Behavioral training

Details of the behavioral task have been described in previous studies (Khan et al., 2018; Poort et al., 2015). Food restricted mice were trained in a virtual environment to perform a visual go-no go discrimination task. Trials were initiated by head-fixed mice running on a Styrofoam wheel for a randomly chosen distance in an approach corridor (black and white circle pattern unrelated to the task for 111cm followed by gray walls for 74-185 cm plus a random distance of gray walls chosen from an exponential distribution with mean 37 cm). Mice were then presented with either a vertical grating pattern (square wave gratings, 100% contrast) or an angled grating pattern (rotated 40° relative to vertical) on the walls of the virtual environment. Mice were rewarded with a drop of soy milk for licking a reward spout in response to the vertical grating (hits). One or more licks in the angled grating corridor were considered errors (false alarms). Mouse performance was quantified using a behavioral d-prime: *bd* ‘ = Φ^−1^ (*H*) − Φ^−1^ (*F*), where Φ ^−1^ is the normal inverse cumulative distribution function, H is the rate of hit trials and F is the rate of false alarm trials.

After reaching high levels of discrimination performance, all mice were trained to switch between blocks of an olfactory and visual discrimination task (the attention switching task). The visual blocks were the same as the visual discrimination task described above. In olfactory blocks, mice performed an olfactory go-no go discrimination task in which odor 1 (10% soya milk odor) was rewarded and odor 2 (10% soya milk with 0.1% limonene mixture) was not rewarded. Odors were delivered through a flow dilution olfactometer calibrated with a mini PID (Aurora) at 10-20% saturated vapor concentration of the above solutions, and at 1 L/min flow rate. Before the presentation of odors, in 70% of randomly chosen trials mice were also presented with the same vertical or angled grating stimuli at different positions in the approach corridor, with random delays from trial start chosen from the same distribution as in the visual block. Mice learnt to ignore these irrelevant grating stimuli while accurately discriminating the odors. On switching to the visual block, mice licked selectively to the rewarded grating as before. Mice typically performed two visual and two olfactory blocks in each session, data was pooled across blocks of the same type. After each block transition, we excluded trials in which the behavior of the mice was ambivalent (Poort et al., 2015). Each block typically contained 70-150 trials. Mice typically learnt to switch successfully within 1-2 days.

### Immunohistochemistry and image registration

Brain fixation was performed by transcardial perfusion with 4 % paraformaldehyde in phosphate buffer 0.1 M followed by 24 hours of post-fixation in the same solution at 4°C. The brains underwent two freeze-thaw cycles in liquid nitrogen, and were sliced tangentially to the surface of visual cortex. 80 µm slices were cut on a vibratome (Zeiss Hydrax V50) and were immunostained for PV, SOM and VIP (Khan et al., 2018). Primary and secondary antibodies are listed in (Khan et al., 2018). We imaged the slices with a confocal microscope (Zeiss LSM 700), and confocal z-stacks were registered with the previously acquired in vivo imaging planes and z-stacks of the recording sites. Cells were identified manually and assigned to cell classes based on immunostaining.

### Data analysis

Regions of interest (ROIs) from motion-corrected image stacks were selected for each cell in each session. We adapted the method of (Chen et al., 2013) to correct for neuropil contamination of calcium traces. Neuropil masks were created for each cell by extending the ROI by 25μm and including all pixels that were more than 10μm away from the cell boundary, excluding pixels assigned to other cells or segments of dendrites and axons (pixels that were more than 2 standard deviations brighter than the mean across all pixels in the neuropil mask). We performed a robust regression on the fluorescence values of the ROI and neuropil mask. We inspected the slope of this regression in a sample of our dataset and obtained a factor of 0.7 by which we multiplied the neuropil mask fluorescence (median subtracted) before subtracting it from the ROI fluorescence to obtain the neuropil-corrected raw fluorescence time series F(t). Baseline fluorescence F_0_(t) was computed by smoothing F(t) (causal moving average of 0.375s) and determining for each time point the minimum value in the preceding 600s time window. The change in fluorescence relative to baseline, ΔF/F, was computed by taking the difference between F and F_0_, and dividing by F_0_. All data used in this study for the learning epoch is the same as that used in (Khan et al., 2018), except with neuropil corrected signals used throughout.

Responses were analyzed for the vertical and angled grating corridor by aligning neuronal activity to the onset of the stimuli. We used a Wilcoxon rank-sum test to determine if the response of a cell (average ΔF/F in a time window of 0-1 s after grating onset) was significantly different between vertical and angled gratings (P < 0.05). We used a Wilcoxon signed-rank test to determine if the response (ΔF/F 0-1 s) to the gratings significantly increased or decreased relative to baseline (−0.5 to 0 s). For visualizing stimulus-evoked responses and for computing the change in stimulus-evoked responses with learning and attention, we subtracted the pre-stimulus baseline (−0.5 to 0 s before stimulus onset) from the average response.

The selectivity of each cell was quantified as the selectivity index (SI), the difference between the mean response (0-1 s) to the vertical and angled grating divided by the pooled standard deviation, which was positive or negative for cells that preferred the vertical or angled grating respectively. We took the average of the absolute selectivity of all cells to obtain an average measure of the selectivity across a population of cells (including vertical and angled preferring cells). We calculated the selectivity of the local PYR population around each PV cell by averaging the responses of all PYR cells, within 100 μm distance, to the two grating stimuli. Confidence intervals were calculated by a bootstrap procedure where we randomly selected cells with replacement 10,000 times to obtain the 2.5 and 97.5 percentiles. The P value was given by the percentage of bootstrapped pre-learning or ignore condition slope values that were lower than the post-learning or attend slope multiplied by two (two-sided test). To compute Δselectivity during learning and attention, we took the difference SI^post^ – SI^pre^ or SI^attend^ – SI^ignore^ for cells with positive selectivity post learning or in the attend condition. Similarly, we took the difference –(SI^post^ – SI^pre^) or –(SI^attend^ – SI^ignore^) for cells with negative selectivity post learning or in the attend condition.

To compute noise correlation, we first subtracted for each trial and each cell the average responses across all trials. We then used the Pearson correlation coefficient to quantify the correlation between responses of pairs of cells. Changes in noise correlations with learning and attention between different cell types were tested using a sign test on all cells imaged pre- and post-learning or in the ignore and attend conditions.

In a previous study based on the learning dataset used here, we controlled for the effects of running and licking on neural responses (Khan et al., 2018, Supplementary figures 5 and 8). Here we performed similar analysis on the attention dataset. We controlled for the possible effect of variations in running speed across the ignore and attend conditions on stimulus selectivity and noise correlations using a stratification approach. We selected a subset of trials with similar distributions of running speed in the ignore and attend condition for each stimulus. We then recomputed the stimulus selectivity and noise correlations in the attend and ignore conditions and obtained similar results with and without stratification (Fig. S3A,C). On excluding trials with licks in the analysis window (0-1 s after grating onset), we also obtained similar results for stimulus selectivity and noise correlations (Fig. S3B,D).

### Linear Multivariate Autoregressive System Model

Details of the MVAR model are described in a previous study (Khan et al., 2018). We fit the activity of all simultaneously imaged neurons using a multivariate autoregressive (MVAR) linear dynamical system incorporating stimulus-related input, the simultaneously measured co-fluctuations from multiple cells of different cell types and the mouse running speed. We estimated the interaction weights between pairs of cells which describe the relationship between the activity of one cell and the activity of another cell at previous timepoints, conditioned over the activity of all other cells and over behavioral and sensory variability.

The learning-related data was previously studied in detail using this model (Khan et al., 2018). Here we fit the model separately to the learning and attention switching tasks, in each case fitting either separate interaction weights for the pre/post learning or ignore/attend conditions or a single set of weights to account for activity in both conditions. The different MVAR models were compared using leave-one-out cross validation (Figure S6B), measuring prediction quality on held-out data. We held out one vertical grating trial from the post learning or attend condition in the test set, using the remaining trials of all types for training. The MVAR model was fit to these training data, and the error in the model prediction was calculated for each time sample in the test trial. This procedure was repeated, leaving out each vertical grating trial in turn. We calculated an *R*^2^ value for each cell combining errors across all of these trials. Specifically, the *R*^2^ was defined relative to a baseline model which incorporated only the trial-averaged response profile of each cell, i.e. *R*^2^ = 1 – (sum of squared errors in MVAR prediction)/(sum of squared errors in the trial-averaged response profile prediction). Running speed was not included in the model for the cross-validation analysis to facilitate comparison with alternative models.

### Circuit model

We modeled a circuit consisting of an excitatory population PYR, and three inhibitory populations, corresponding to PV, SOM, and VIP interneurons. The activity of the population *i* is described by its calcium response *r*_*i*_, which evolves over time according to one of the following equations:

Additive model:

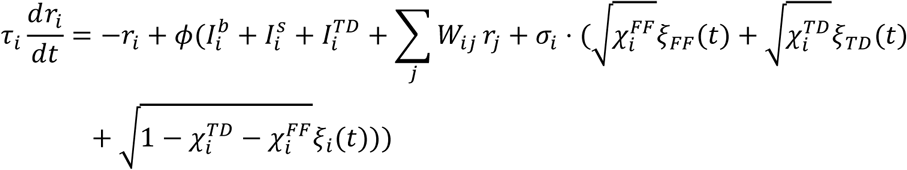

Multiplicative model:

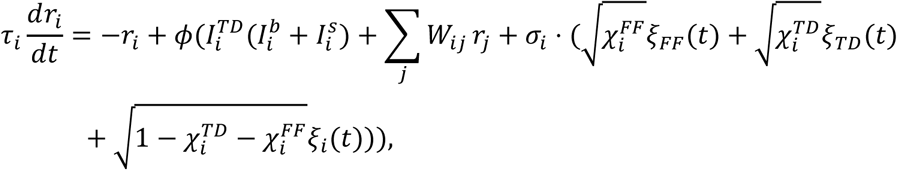

where *i, j* ∈ {*PYR, PV, SOM, VIP*} and

*τ*_*i*_ is the time constant of population *i*.

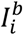 is the baseline input to population *i*,

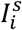 is the stimulus-dependent feedforward input to population *i*,

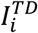 is the modulatory top-down input - the attentional modulation of population *i*, and

∑_*i*_ *W*_*ij*_*r*_*j*_ is the recurrent input from the local circuit and *W*_*ij*_ is the effective synaptic weight. As in earlier models (Kanashiro et al., 2017), each population received private and shared noise. *ξ*_*i*_(*t*) is noise, private to each population, corresponding to noise arising from ion channels, or the activation function.

*ξ*_*TD*_(*t*) and *ξ*_*FF*_(*t*) are shared noise terms arising from shared modulatory top-down and/or feedforward inputs. *ξ*_*i*_(t), *ξ*_*TD*_(*t*), and *ξ*_*FF*_(*t*) are drawn from a Gaussian distribution with zero mean and unit variance. We assume that external noise sources contribute equally. *ϕ*(*x*) is the activation function:

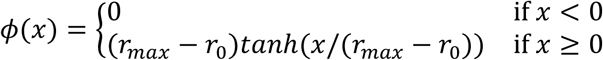

PYR and PV populations receive an input current 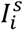 upon presentation of their preferred stimulus (Ji et al., 2016) representing thalamic inputs. They receive a fraction of this input current (0.2.*I*_*s*_) upon presentation of their non-preferred stimulus. Similar results were observed when SOM and VIP populations also received the same input current as PV cells. All populations received a constant baseline current input 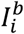. Each modulated population *i* received a top-down modulation 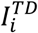, which took one of two values {*x*_*ignore*_, *x*_*attend*_} depending on the absence or presence of attention (see Tables 1 and 2). *r*_0_ = 1.0 and *r*_*max*_ = 20.0 denote the minimum and maximum activity, respectively.

**Table 1:** Inputs to the multiplicative model. Shown are the values for the baseline, stimulus, and top-down inputs to the populations PYR, PV, SOM, and VIP. Top-down inputs depend on the condition, which is either ignore or attend: {*x*_*ignore*_, *x*_*attend*_}.

| Population | baseline $I_i^b$ | stimulus $I_i^s$ | top-down $I_i^{TD}$ |
| --- | --- | --- | --- |
| PYR | 6.0 | 17.8 | {1.0, 2.0} |
| PV | 4.0 | 10.0 | {1.0, 2.0} |
| SOM | 1.2 | 0.0 | {1.0, 2.0} |
| VIP | 4.6 | 0.0 | {1.0, 2.0} |

**Table 2:** Inputs to the additive model. Shown are the values for the baseline, stimulus, and top-down inputs to the populations PYR, PV, SOM, and VIP. Top-down inputs depend on the condition, which is either ignore or attend: {*x*_*ignore*_, *x*_*attend*_}.

| Population | baseline $I_i^b$ | stimulus $I_i^s$ | top-down $I_i^{TD}$ |
| --- | --- | --- | --- |
| PYR | 6.0 | 17.8 | {0.0, 1.0} |
| PV | 4.0 | 10.0 | {0.0, 1.0} |
| SOM | 1.2 | 0.0 | {0.0, 1.0} |
| VIP | 4.6 | 0.0 | {0.0, 1.0} |

We changed the contributions of noise sources to the overall noise in the populations, depending on the inputs population *i* received, according to Kanashiro et al. (Kanashiro et al., 2017). If population *i* received attentional modulation:

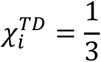

otherwise:

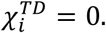

If population *i* received feedforward input:

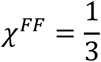

otherwise:

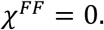

The standard deviation of the total noise was given by:

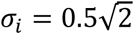

## Connectivity

We took the weight matrix *W* from (Kuchibhotla et al., 2017), and adjusted only the baseline and stimulus inputs 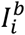 and 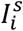 such that the simulated neural responses matched the data.

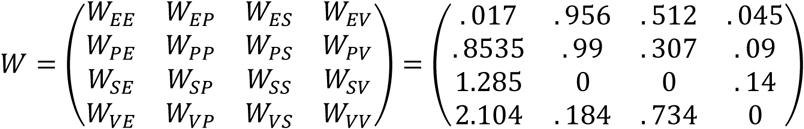

Each population was represented twice in the model, allowing us to measure noise correlations within cell classes.

We simulated the network without stimulus input for 5s until the neural activity for each cell class reached steady state. Then we presented the non-preferred stimulus for 3s, following which we waited another 4s before we presented the preferred stimulus for 3s. The simulation time step was 1ms. We repeated this protocol for 100 trials. *τ*_*PYR*_ was 800ms and *τ*_*i*_ with *i* ∈ {*SOM, VIP, PV*} was 400ms.

To calculate the selectivity of cell populations in the model, we subtracted the mean activity to the non-preferred stimulus 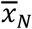 from the mean activity to the preferred stimulus 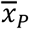 during 1s after stimulus onset and normalized by their pooled standard deviation *s*_*pooled*_:

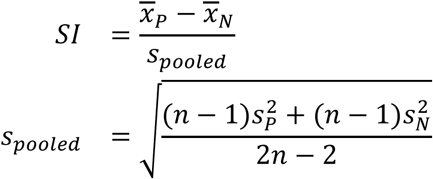

where *n* is the number of trials, *s*_*P*_ is the standard deviation of the activity during the preferred stimulus, and *s*_*N*_ is the standard deviation of the activity during the non-preferred stimulus.

To determine the noise correlation between cell populations in the model, we calculated the average activity in populations *x* and *y* in each trial *i* in a 1s time window after onset of the preferred stimulus: *x*_*i*_ and *y*_*i*_. We calculated the means 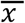 and 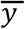 and standard deviations *σ*_*x*_ and *σ*_*y*_ of the activity over trials for each population. We then calculated noise correlations between populations *x* and *y* over *n* = 100 trials according to the following equation:

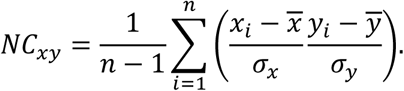

For Figure S6D, 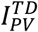 and 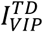 were 0.0, and we varied 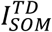 continuously between 1 and 2.2 and 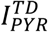 proportionally to 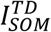 as indicated in the figure.

