## Supplementary Figures for "Learning and attention increase visual response selectivity through distinct mechanisms"

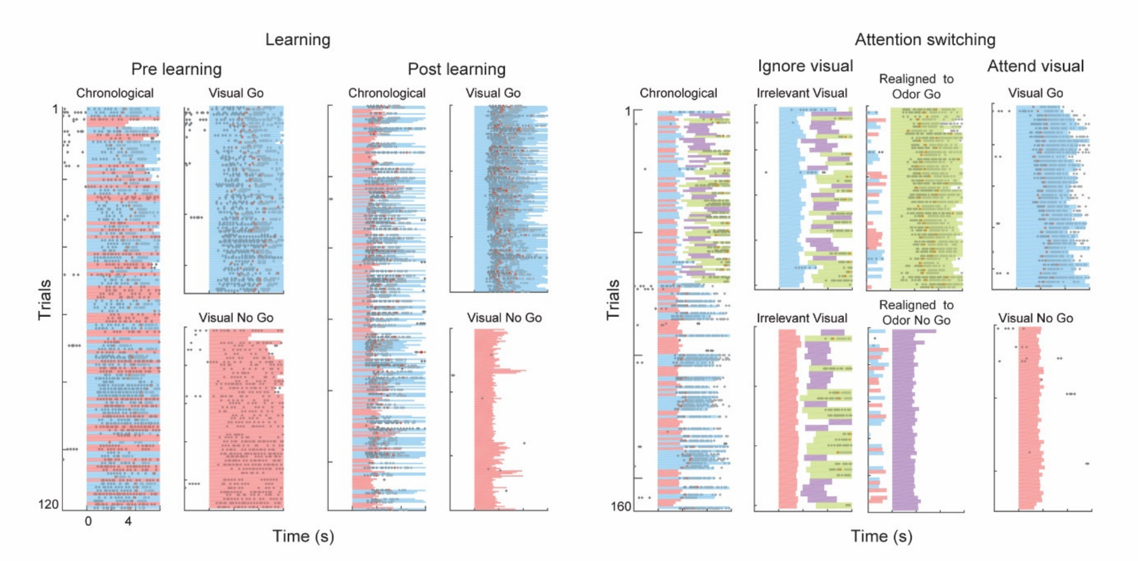


**Figure S1. Example behavior sessions.** Left, lick rasters from example sessions pre- and post-learning. Right, example session of attention switching task, one block each of ignore and attend visual stimuli. Each row is a trial aligned to stimulus onset, black dots indicate licks, red dots indicate reward delivery, red and blue shading indicates presence of vertical and angled visual grating stimuli respectively, green and purple indicates odor1 and odor2 delivery respectively.

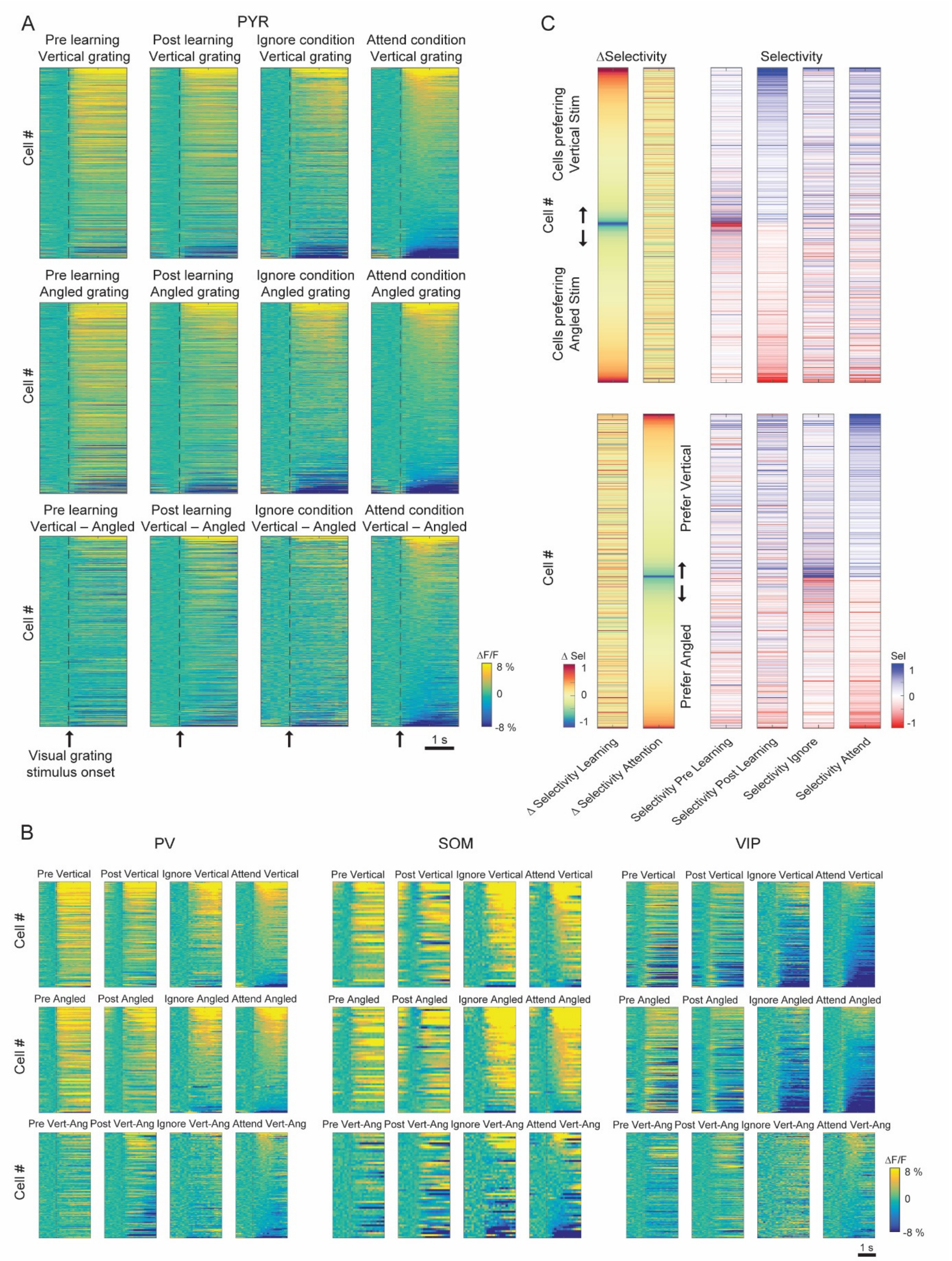


**Figure S2. Responses and selectivity of cells across learning and attention switching.** A) Average responses of all PYR cells that were matched across the learning and attention tasks (N = 915 cells). Responses are shown pre and post learning and in the ignore and attend conditions (columns). Responses are aligned to the vertical grating, angled grating and the difference between the two (rows). Cells are sorted in the final column (attend condition) by their average response amplitude 0–1 s from stimulus onset, and the remaining three panels in the same row are shown with the same cell sorting, to aid comparing the same cells' responses in different conditions. All responses are baseline corrected (subtraction of baseline Δ F/F –0.5 to 0 s before stimulus onset) and aligned to grating onset (dashed line). B) Same as A) for the three interneuron classes, N = 105 PV cells, 54 SOM cells and 144 VIP cells. C) ΔSelectivity for the same cells during learning and attention displayed in color code (left, similar to Figure 3A). The same cell sorting is maintained throughout to show the selectivity of the same cells in the different conditions (right). Top and bottom are the same data sorted differently; cells are sorted by Δselectivity during learing (top) or attention (bottom), and by splitting the data into those cells which prefer vertical or angled stimuli in the post learning or attend condition respectively (indicated by arrows).

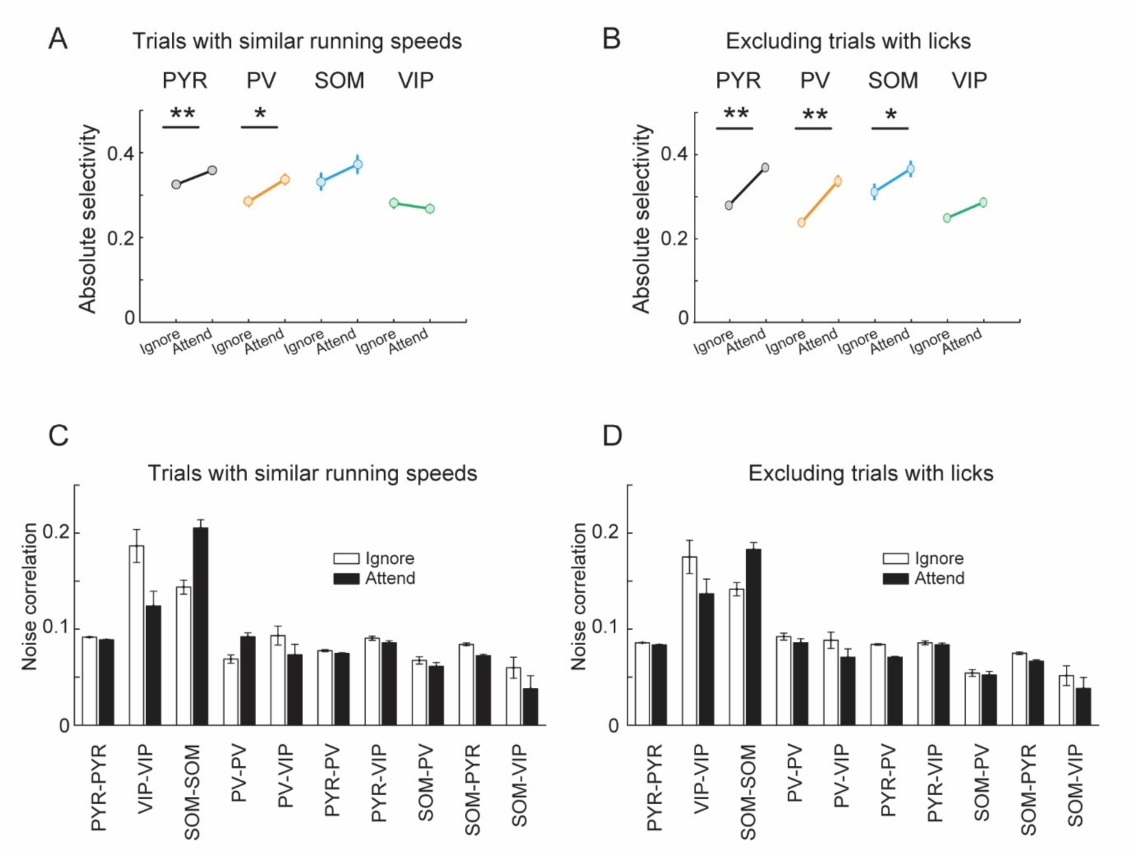


**Figure S3. Differences in running speed and licking cannot account for the pattern of changes in stimulus selectivity and noise correlations.** A) Mean absolute selectivity of each cell class in the ignore and attend conditions (computed in the period of 0-1s after grating onset) after equalizing the distributions of running speed in the two conditions for each stimulus presentation. B) Mean absolute selectivity of each cell class when excluding all trials with licks. Sign test, **, P < 0.001; *, P < 0.05. C-D) Similar analysis for noise correlations measured during the vertical grating response (0-1 s from stimulus onset). Error bars represent SEM. Similar analysis was done on the learning dataset in Khan et al 2018 showing that changes in running speed and licking could not account for the pattern of changes in stimulus selectivity and noise correlations during learning.

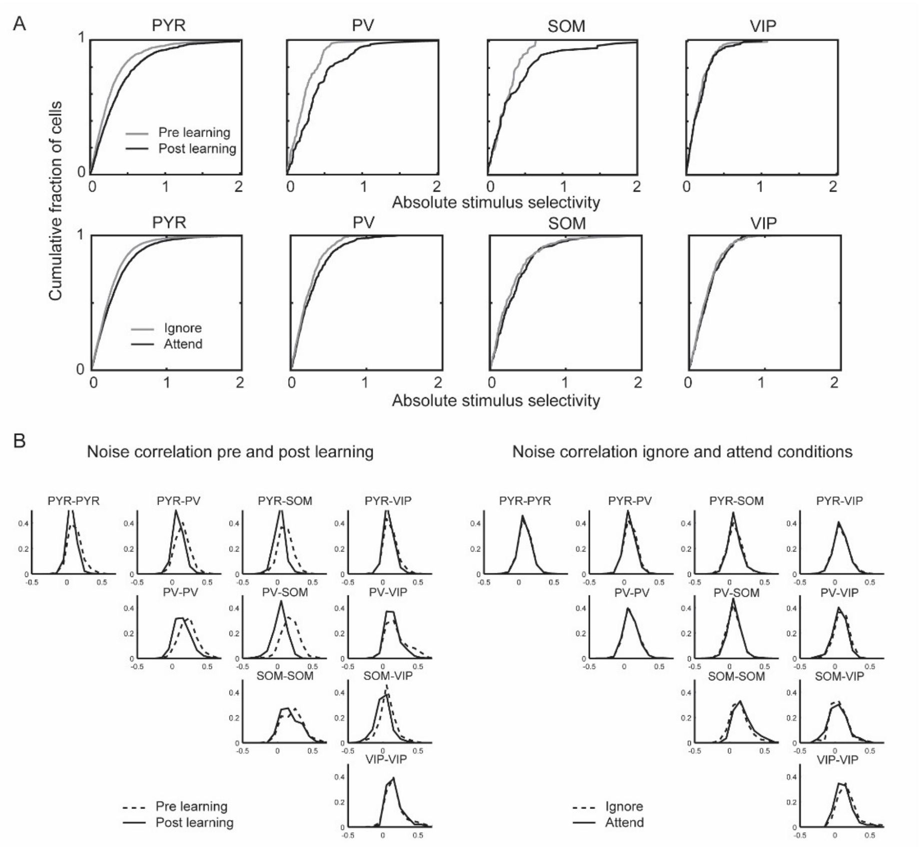


**Figure S4. Distributions of selectivity and noise correlation.** A) Cumulative histograms of stimulus selectivity of each cell class. Selectivity was measured during the grating response (0-1 s from stimulus onset) before and after learning, and in the ignore and attend condition of the attention switching task. The number of cells pre, post-learning was: 1,249 PYR, 132 PV, 58 SOM and 175 VIP cells; ignore, attend condition: 5813 PYR, 477 PV, 245 SOM and 365 VIP cells. B) Distributions of noise correlation between cell pairs of each combination of cell classes during the vertical grating stimulus presentation. Noise correlation was measured during the grating response (0-1 s from stimulus onset) between cell pairs of each combination of cell classes, before and after learning (left), and in the ignore and attend condition of the attention switching task (right). The number of cell pairs in each cell class combination was: pre-, post-learning, PYR–PYR 74,581, 64,921; VIP–VIP 1166, 907; SOM–SOM 215, 99; PV–PV 1,731, 1,369; PV–VIP 790, 718; PV–PYR 17,792, 15,283; PYR–VIP 14,681, 12,009; SOM–PV 1,250, 690; SOM–PYR 7,112, 4,952; SOM–VIP 455, 377. Ignore/attend conditions, PYR–PYR 61,175; VIP–VIP 58; SOM–SOM 381; PV–PV 777; PV–VIP 129; PV–PYR 11,312; PYR–VIP 3024; SOM–PV 814; SOM–PYR 6,626; SOM–VIP 136.

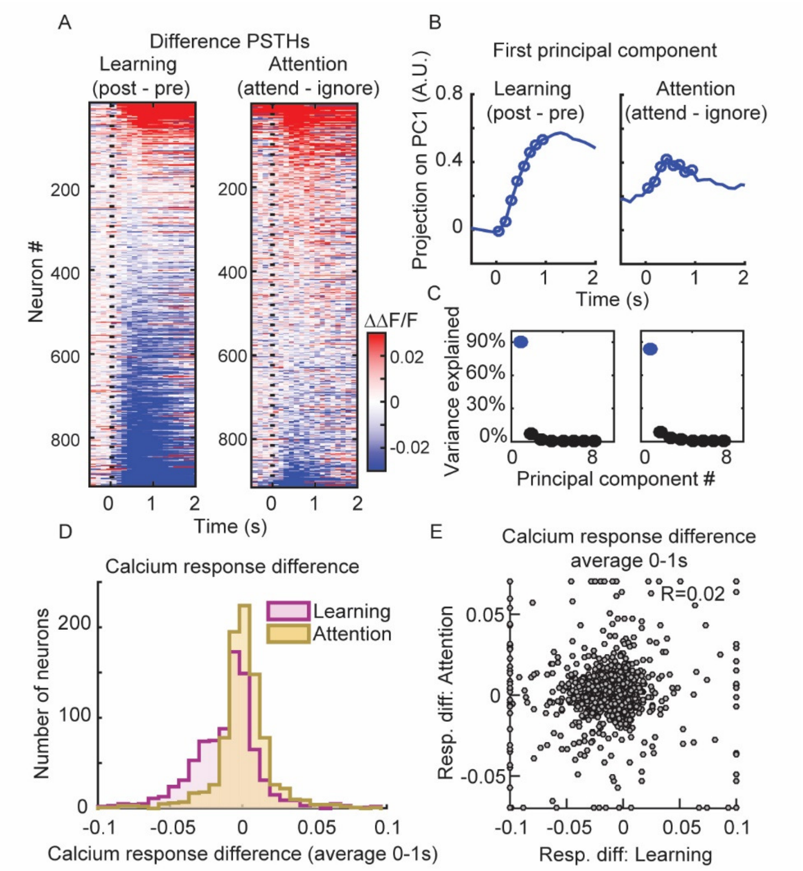


**Figure S5. Response changes during learning attention**. A-C) Similar to Figure 4A-C for non-rewarded angled stimulus. A) Difference in calcium responses to the non-rewarded angled stimulus, post minus pre learning (left) or attend minus ignore conditions (right) for all matched PYR cells (Difference-PSTHs). Responses are baseline corrected (subtraction of baseline ΔF/F –0.5 to 0 s before stimulus onset) and aligned to grating onset (dashed line). Cells are sorted by their average amplitude 0–1 s from stimulus onset. N = 915 matched cells here and below. B) First principal component (PC) of the difference-PSTHs from the learning (left) and attention data (right). Circles indicate the time points (0-1s) used to determine the PCs. C) Percentage of variance explained by each PC during learning (left) and attention (right). D) Distribution of average calcium response difference (difference-PSTHs averaged 0-1s) in response to rewarded vertical grating stimulus, during learning and attention ($P < 10^{-28}$, sign test). E) Relationship between the calcium response difference during learning and attention ($R = 0.02$, $P = 0.51$). Values greater than the axis limits are pegged to the maximum displayed value. Similar results as D and E were obtained with non-rewarded angled gratings, data not shown.

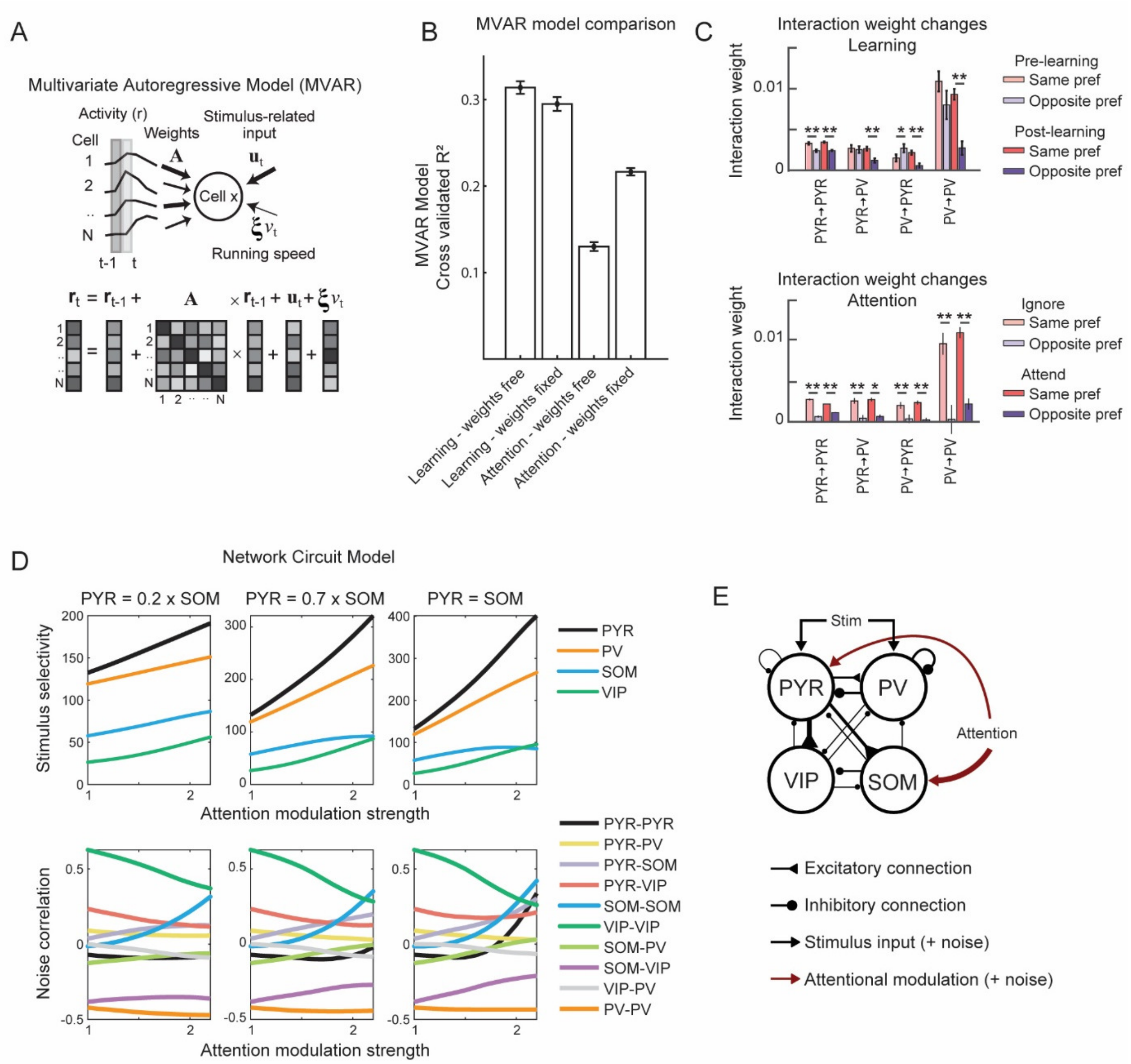


**Figure S6. Computational modelling of learning and attention-related activity changes.** A) Schematic depicting the MVAR model which fits single-trial responses by estimating the contribution of stimulus-locked input, recurrent inputs from the local cell population and running speed. B) Comparison of different MVAR models. Cross-validated $R^2$ of different versions of the MVAR model fit to data with different constraints. When fitting pre- and post-learning data, cross-validated $R^2$ is higher when interaction weights are allowed to change from pre to post learning (learning: weights free, learning: weights fixed). When fitting attention data, cross-validated $R^2$ is lower when interaction weights are allowed to change between ignore and attend conditions (attention: weights free, attention: weights fixed). C) In an MVAR model where weights were allowed to change, average interaction weights are shown for cell pairs of specific cell classes, and with the same or opposite stimulus-input preference before and after learning (top) or during ignore and attend conditions (bottom). Error bars indicate SEM. Stronger weights between same orientation preference pairs emerged during learning, and this pattern did not change with attention. D) Changes in stimulus selectivity (top) and noise correlation between cells (bottom) for varying degrees of attention modulation applied to SOM and PYR cells. Three combinations are shown with varying degrees of modulation applied to PYR relative to SOM populations. Left: model with PYR = 0.2×SOM modulation. Middle: model with PYR = 0.7×SOM modulation. Right: model with PYR = SOM modulation. Modulation of PYR and SOM populations with PYR = 0.7×SOM modulation best fits the data. C) Schematic showing the final circuit model which best accounts for the data.
